# Immunometabolic Reprogramming by Thyroid-Stimulating Antibodies Drives Orbital Adipogenesis via Histone Lactylation in Thyroid Eye Disease

**DOI:** 10.64898/2026.07.27.741014

**Authors:** Yanchen Zhang, Neng Ling, Min Wu, Ling Zeng, Ben Chen, Weibin Liu, Zhixing Li, Yan Zhang, Xiaojing Li, Xuanwen Li, Yicheng Mao, Yate Huang, Shiguan Zhou, Shuhui Wan, Hongmei Wang, Yunhai Tu, Jianzhang Wu, Mao Ye, Wencan Wu

## Abstract

Thyroid eye disease (TED) is conventionally viewed as an autoimmune inflammatory disorder. However, the metabolic determinants driving orbital tissue remodeling remain largely undefined. Here, we uncover a paradigm-shifting mechanism whereby thyroid-stimulating antibodies (TSAbs) instigate profound metabolic reprogramming in orbital fibroblasts (OFs), driving a switch toward glycolysis and markedly elevating lactate production via the CREB/LDHA axis. Critically, we demonstrate that this TSAbs-induced metabolic perturbation is not a mere byproduct but the principal driver of pathology. Mechanistically, lactate functions as a signaling metabolite that directly potentiates adipogenesis via histone lactylation-dependent transcriptional activation of the master adipogenic regulators PPARγ and C/EBPα. Collectively, our findings underscore metabolic reprogramming as a central pathogenic mechanism in TED, whereby autoimmune cues drive disease progression through metabolic-epigenetic crosstalk. This work substantiates the paradigm of TED as an immune-metabolic disorder.

## 1. Introduction

Thyroid eye disease (TED) is an organ-specific autoimmune disorder characterized by proptosis, eyelid retraction, and restrictive strabismus, often leading to disfigurement, vision impairment, and reduced quality of life^1^. Pathologically, TED is driven by the autoimmune-mediated aberrant activation of orbital fibroblasts (OFs), which leads to inflammatory infiltration, adipose tissue hyperplasia, and fibrotic remodeling^1, 2^. Although the immunological features of TED have been extensively characterized, how autoimmune signaling is coupled to the metabolic reprogramming that drives tissue remodeling remains poorly understood. OFs serve as both key effector cells and targets of autoimmune attack, undergoing profound metabolic changes that direct their differentiation into myofibroblasts and adipocytes. Thyroid-stimulating antibodies (TSAbs) targeting the thyrotropin receptor (TSHR) mediate a cross-reactive immune response that extends beyond thyroid cells to TSHR-expressing OFs, adipocytes, and extraocular muscle cells^1^. Consequently, serum TSAbs levels strongly correlate with the clinical activity and severity of the disease^3–5^, yet the mechanisms that transduce TSHR signaling into the metabolic reprogramming of OFs have not been defined.

Adipose tissue expansion is a central pathological feature of TED. Orbital adipose tissues (OATs) from patients with TED contain an increased number of smaller adipocytes compared to those in controls, suggesting that adipogenesis is the primary driver of tissue enlargement^6^. Under these pathological conditions, OFs differentiate into adipocytes through a process orchestrated by the master transcription factors PPARγ and C/EBPα^7^. Therefore, understanding how extracellular signals, particularly TSAb-mediated TSHR activation, engage these transcriptional programs is a central question in TED pathogenesis.

Metabolic reprogramming, the systematic rewiring of cellular energy and biosynthetic metabolism, is a hallmark of diverse pathological conditions, including tumorigenesis, neurological disease and immune dysregulation^8^. In TED, the increased bioenergetic and biosynthetic demands of orbital tissues are met by upregulated glycolysis^9–11^. However, whether this metabolic shift actively drives disease progression remains unclear. Lactate, the terminal metabolite of glycolysis, was long regarded as a metabolic byproduct but is now recognized as a signaling molecule with diverse cellular functions, including immune regulation, metabolic adaptation, and tissue regeneration^12–14^. Lactate also serves as a substrate for protein post-translational modification, lysine lactylation, which regulates gene expression in a lactate concentration-dependent manner^15^. Lactylation has been implicated in chromatin remodeling, gene expression control, and cell fate determination^12, 14^. Although elevated glycolysis has been reported in TED^10^, the specific contributions of lactate and lactylation to adipose hyperplasia, as well as their mechanistic links to TSHR signaling, remain unknown.

In this study, we identify a TSHR-linked immunometabolic-epigenetic cascade in TED. We demonstrate that lactate and lactylation serve as promising biomarkers for the disease. Mechanistically, TSAbs drive metabolic reprogramming by upregulating lactate production via the CREB-LDHA-lactate axis, which in turn promotes adipogenesis through H3K18 lactylation.

## 2. Results

### 2.1. Lactic acid levels are elevated in TED and positively correlated with serum TSAb levels

To determine whether glycolysis is dysregulated in TED, we reanalyzed a public single-cell RNA-seq dataset of human orbital adipose tissue from the GEO database (GSE194323). The t-SNE analysis identified orbital fibroblasts (OFs) as the largest cell population, accounting for nearly half of all cells in this tissue (Fig. 1A). We next generated a heatmap of glycolysis-related genes expressed across orbital tissues using public scRNA-seq data from normal donors and patients with TED. Rate-limiting glycolytic enzymes (PKM, PFK) and the lactate-producing enzyme lactate dehydrogenase A (LDHA) were significantly upregulated in multiple cell populations of orbital adipose tissue in TED, most notably in OFs (Fig. 1B). To directly assess metabolic remodeling in TED, we collected OATs from patients with TED and normal donors and performed targeted energy metabolomics profiling. Principal component analysis (PCA) score plots revealed a clear separation between the metabolic profiles of OATs from TED patients and those from normal donors (Fig. 1C). Metabolite set enrichment analysis (MSEA) further identified the glycolysis/gluconeogenesis pathway (KEGG hsa00010) as significantly enriched (Fig. 1D). Consistent with this enrichment, key glycolysis-associated metabolites, such as lactate, were markedly altered, showing a robust and consistent increase in TED (Fig. 1E). We confirmed this by quantifying lactate concentrations in orbital samples using a colorimetric assay, which showed that lactate levels were significantly elevated in samples from both TED patients and TED mice (Fig. 1F, M), consistent with our targeted metabolomic analysis. Receiver operating characteristic (ROC) analysis revealed that orbital lactate levels exhibited excellent diagnostic performance for TED (AUC = 0.9188) (Fig. 1G). Furthermore, correlation analysis showed that lactate levels in human OATs were positively correlated with serum TSAb levels (R² = 0.7533), Clinical Activity Score (CAS) (R² = 0.8359), and proptosis (R² = 0.8122) (Fig. 1H-J). Similarly, lactate levels in mouse orbits also exhibited a robust positive correlation with serum TSAb levels (R² = 0.9071) (Fig. 1N). Taken together, these results demonstrate that lactate is a significantly upregulated biomarker in TED that tracks with disease severity, implicating lactate metabolism in TED progression.

**Figure 1.**
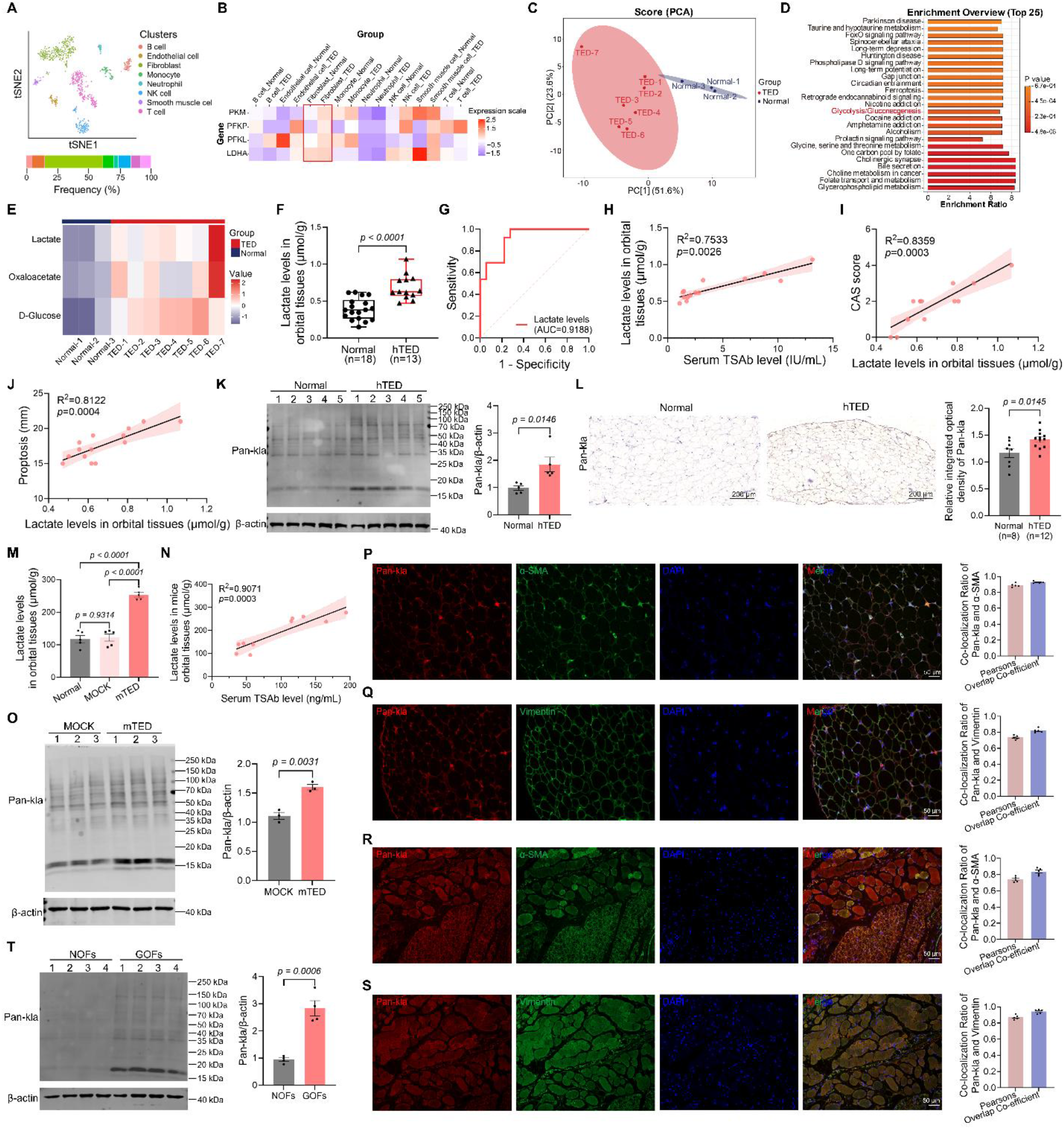
Lactic acid levels are elevated in TED and positively correlated with serum TSAb levels. (A) Reanalysis of a public single-cell RNA-seq dataset from human orbital adipose tissues. The t-SNE map shows different cell types in human orbital adipose tissues. (B) Heatmap showing the expression of four glycolysis-related genes in orbital tissues from normal donors and TED patients, based on publicly available single-cell RNA sequencing data. (C) Principal component analysis (PCA) score plots of metabolic profiles showing metabolomic variation among orbital tissues from normal donors (n = 3) and TED patients (n = 7). (D) Bar plot showing the top 25 metabolic pathways enriched in the metabolite set enrichment analysis (MSEA). (E) Heatmap of glycolytic metabolite levels. (F) Lactate levels of orbital adipose tissues in normal donors (n = 18) and TED patients (n = 13). (G) Receiver operating characteristic (ROC) analysis evaluating the diagnostic performance of orbital lactate levels for TED. AUC, area under the curve. (H) The correlation between lactate levels in orbital tissues and serum TSAb levels was evaluated using Spearman correlation analysis, with the *p*-value determined by permutation test. (I) The correlation between the clinical activity score (CAS) and lactate levels in orbital adipose tissues of TED patients was evaluated using Spearman correlation analysis, with *p*-values determined by a permutation test. (J) The correlation between the degree of proptosis and lactate levels in orbital adipose tissues of TED patients was evaluated using Pearson correlation analysis, with *p*-values determined by Student’s t-tests. (K) The global lactylation levels of orbital adipose tissues in normal donors (n = 5) and TED patients (n = 5). β-actin served as the loading control. (L) IHC staining images of Pan-kla in orbital adipose tissues from normal donors (n = 8) and TED patients (n = 12) and corresponding quantitative analysis of relative integrated optical density. Scale bar = 200 μm. (M) Lactate levels were measured in the orbital tissues of mice from the Normal, MOCK, and mTED groups (n = 5). (N) The correlation between the lactate levels in mouse orbital tissues and serum TSAb levels was evaluated using Pearson correlation analysis, with *p*-values determined by Student’s t-tests. (O) The global lactylation levels of orbital tissues in MOCK mice (n = 3) and TED mice (n = 3). β-actin served as the loading control. (P-Q) Representative immunofluorescence images of Pan-kla (red) and α-SMA or Vimentin (green) in orbital adipose tissues from TED patients (left panel, scale bar = 50 μm). Colocalization analysis using Pearson’s correlation coefficient and the overlap coefficient of Pan-kla and α-SMA or Vimentin in orbital adipose tissues (right panel, n = 5). (R-S) Representative immunofluorescence images of Pan-kla (red) and α-SMA or Vimentin (green) in orbital adipose tissues from TED mice (left panel, scale bar = 50 μm). Colocalization analysis using Pearson’s correlation coefficient and the overlap coefficient of Pan-kla and α-SMA or Vimentin in orbital adipose tissues (right panel, n = 5). (T) The global lactylation levels of OFs from normal donors (n = 4) and TED patients (n = 4). β-actin served as the loading control. All data are represented as the mean ± SEM. Two-tailed unpaired Student’s t-tests were performed in (F, K, L, O, T). One-way ANOVA, followed by Tukey’s multiple post hoc test, was performed in (M). Accurate *p*-values are listed in the figures.

Because lactate can serve as a direct substrate for lactylation^15^, we examined whether elevated lactate levels promoted protein lactylation in orbital tissues. To characterize the lactylation landscape in TED, we assessed global lactylation levels using western blotting and immunohistochemistry (IHC) with an anti-pan-lactylation antibody. Global lactylation was significantly elevated in orbital tissues from TED patients and TED mice compared with normal donors and healthy mice (Fig. 1K-L, O). To determine the cellular localization of lactylated proteins, we performed immunofluorescence co-staining of orbital tissues using the fibroblast markers α-SMA and Vimentin. The results revealed extensive colocalization, with both Pearson’s correlation coefficients and overlap coefficients exceeding 0.75 (Fig. 1P-S). To extend these tissue-level findings, OFs were isolated from TED patients (Graves’ orbital fibroblasts, GOFs) and healthy donors (normal orbital fibroblasts, NOFs). Western blot analysis confirmed that global lactylation levels were significantly higher in GOFs than in NOFs (Fig. 1T). Taken together, these data indicate a coordinated elevation of lactate and protein lactylation in TED orbital tissues, implicating the lactate-lactylation axis in TED pathogenesis.

### 2.2. Lactate promotes adipogenesis in vitro and in vivo

Lactate, the terminal metabolite of glycolysis, can function as a signaling metabolite. To minimize potential confounding effects from extracellular pH, we treated GOFs with 5 mM and 10 mM sodium lactate (Nala) to mimic the elevated lactate levels observed in TED orbital tissues. Nala treatment enhanced adipogenic differentiation, as shown by a rounded morphology and increased accumulation of red-stained lipid droplets in a concentration-dependent manner (Fig. 2A). Quantification of cellular triglycerides and BODIPY-stained fatty acids showed increased lipid accumulation in a concentration-dependent manner (Fig. 2B-D). Adipogenesis relies on pro-adipogenic transcription factors, with PPARγ and C/EBPα serving as key regulators^7^. Upon Nala treatment, the protein levels of PPARγ and C/EBPα were increased (Fig. 2E-F).

**Figure 2.**
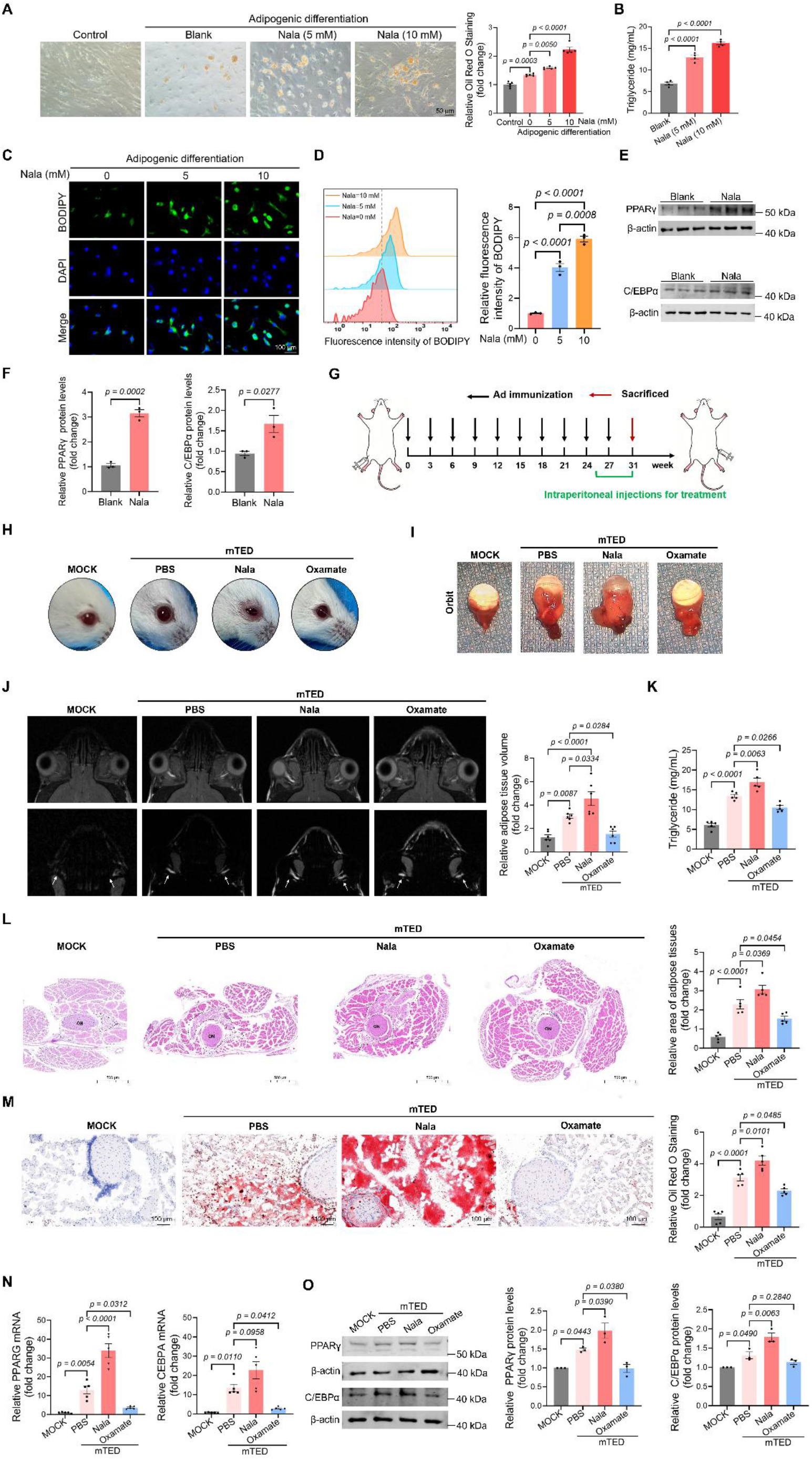
Lactate promotes adipogenesis in vitro and in vivo. (A) Representative Oil Red O staining images of OFs following adipogenic differentiation and Nala treatment. Scale bar = 50 μm. (B) Triglyceride content in adipogenic-differentiated OFs following Nala treatment (n = 3). (C) Representative BODIPY staining images of adipogenic-differentiated OFs following Nala treatment. Scale bar = 100 μm. (D) Flow cytometry analysis of adipogenic-differentiated OFs following Nala treatment (n = 3). (E) Western blot analysis of PPARγ and C/EBPα in adipogenic-differentiated OFs following Nala treatment (n = 3). β-actin served as the loading control. (F) Corresponding quantitative analysis of relative PPARγ and C/EBPα protein levels in panel (E). (G) Experimental timeline for model construction and treatment of the mice. (H) Representative images of the ocular surfaces of mice in the four experimental groups. (I) Representative images of the orbits of mice in the four experimental groups. (J) Representative images showing T2-weighted MRI (upper panels) and fat-water separation maps (lower panels) of mice in the four experimental groups. Corresponding quantitative analysis of the relative adipose tissue volume based on MRI (n = 6). (K) Triglyceride content in the orbits of the four experimental groups. (L) Representative Hematoxylin-eosin (H&E) staining images of orbital tissue sections from the four mouse groups and the corresponding quantitative analysis of the relative area of adipose tissues (n = 5). ON, optic nerve. The white arrow indicates the adipose tissue. Scale bar=500 μm. (M) Representative Oil Red O staining images of orbital tissue sections from the four mouse groups (n = 5). Scale bar = 100 μm. (N) The mRNA expression levels of PPARG and CEBPA in orbital tissues from the four mouse groups by qRT-PCR (n = 5). (O) Western blot analysis of PPARγ and C/EBPα protein levels in orbital tissues from the four mouse groups (n = 3). β-actin served as the loading control. All data are represented as the mean ± SEM. Two-tailed unpaired Student’s t-tests were performed in (F). One-way ANOVA, followed by Tukey’s multiple post hoc test, was performed in (A, B, D, J-O). Accurate *p*-values are listed in the figures.

To test whether lactate contributes to adipose remodeling in vivo, we established a mouse TED model based on a previous report^16^. In this model, Nala was administered to increase orbital lactate levels, whereas Oxamate (an LDHA inhibitor) was used to reduce them. As illustrated in the experimental timeline (Fig. 2G), after nine adenoviral injections to establish the immune induction model, mice with TED were randomly assigned to three treatment groups: intraperitoneal injection with PBS (mTED+PBS), Nala (mTED+Nala), or Oxamate (mTED+Oxamate). To confirm the effectiveness of the intervention, lactate levels in orbital tissues were measured across the treatment groups. The mTED+Nala group showed increased lactate levels compared with mTED+PBS group, whereas the mTED+Oxamate group exhibited reduced lactate levels. We further assessed pan-lactylation levels in orbital tissues by IHC and western blotting. Consistent with lactate measurements, pan-lactylation levels were elevated in the mTED+Nala group and decreased in the mTED+Oxamate group relative to the mTED+PBS control (Supplementary Fig. 1A-C). These findings validate the pharmacologic modulation of the lactate-lactylation axis.

Phenotypically, mice in the mTED+Nala group developed prominent signs of inflammation, including eyelid swelling and congestion, which closely resembled the symptoms observed in TED patients. In contrast, these clinical manifestations were significantly alleviated in the mTED+Oxamate group (Fig. 2H). Orbital imaging showed that Nala treatment promoted orbital tissue overgrowth, which was substantially reduced in the mTED+Oxamate group (Fig. 2I). To quantify adipose content, we performed T2-weighted magnetic resonance imaging (MRI) with water-fat separation. The results revealed significantly increased adipose tissue volume in the orbits of the mTED+Nala group, whereas this pathological adipogenesis was markedly alleviated in the mTED+Oxamate group (Fig. 2J). Histological analysis using H&E and Oil Red O staining showed an expanded adipose area in the mTED+Nala group, which was markedly attenuated in the mTED+Oxamate group (Fig. 2L-M). In parallel, both the mRNA and protein levels of the key adipogenic transcription factors PPARγ and C/EBPα were upregulated in the mTED+Nala group and significantly downregulated in the mTED+Oxamate group (Fig. 2N-O). Overall, these results support that lactate promotes adipogenesis in TED, both in vivo and in vitro.

### 2.3. TSAbs drive lactate production and protein lactylation via CREB-mediated transcriptional activation of LDHA

As noted above, lactate and pan-lactylation in TED orbital tissues correlate with serum TSAb levels. Given that lactate is a byproduct of glycolysis, we investigated the role of TSAb in this metabolic pathway. To this end, we utilized M22 (a human thyroid-stimulating antibody) to perform an extracellular acidification rate (ECAR) assay. M22 treatment enhanced the glycolytic stress response in OFs (Fig. 3A). Given that LDHA catalyzes lactate synthesis, we examined whether M22 regulates LDHA expression. M22 significantly upregulated LDHA at both the mRNA and protein levels (Fig. 3C-D). Consistently, LDHA mRNA and protein levels were significantly elevated in orbital tissues from TED patients and TED mice compared to normal donors and controls (Supplementary Fig. 2A-D). Lactate dehydrogenase B (LDHB) catalyzes the conversion of lactate to pyruvate^17^. In contrast, M22 treatment had no significant impact on LDHB mRNA or protein levels (Supplementary Fig. 3A-B). We next measured intracellular lactate levels in OFs and found that M22 promoted lactate accumulation in a time-dependent manner (Fig. 3E). Consistently, global lactylation levels in OFs increased upon M22 treatment (Fig. 3F). To determine whether these effects depend on LDHA, we used Oxamate, an LDHA-specific inhibitor. Oxamate treatment reversed M22-induced lactate production and global protein lactylation (Supplementary Fig. 4A-B). To rule out off-target effects of pharmacological intervention, we further used siRNA to knock down LDHA expression. All three si-LDHA sequences effectively reduced LDHA at both the mRNA and protein levels (Supplementary Fig. 5A-B). Consistently, si-LDHA reversed M22-induced lactate production and global protein lactylation (Fig. 3G-H).

**Figure 3.**
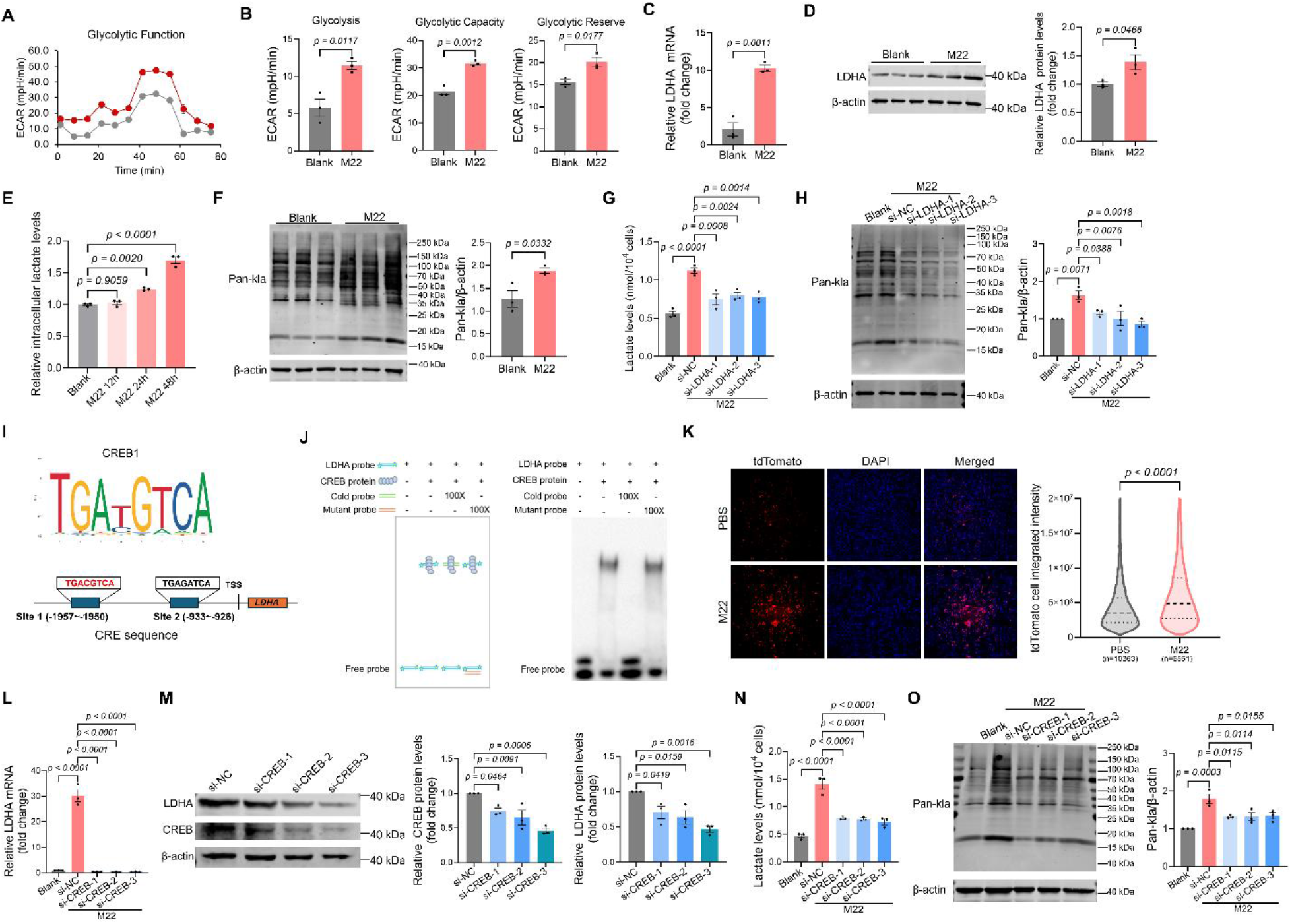
TSAbs drive lactate production and protein lactylation via CREB-mediated transcriptional activation of LDHA. (A) The glycolytic stress response of OFs, with or without M22 treatment, was monitored in real time using the Seahorse system (n = 3). (B) Extracellular acidification rate (ECAR) was used to assess cellular glycolysis (left panel, n=3), glycolytic capacity (middle panel, n=3) and glycolysis reserve (right panel, n=3). (C) The mRNA expression levels of LDHA in OFs stimulated with or without M22 by qRT-PCR (n = 3). (D) Western blot analysis of LDHA expression in OFs stimulated with or without M22 (n = 3). β-actin served as the loading control. (E) Relative intracellular lactate levels in M22-stimulated OFs for the indicated durations (n = 3). (F) Western blot analysis of the global lactylation levels of OFs stimulated with or without M22 (n = 3). β-actin served as the loading control. (G) Lactate levels in OFs transfected with the indicated siRNAs and stimulated with or without M22 (n = 3). (H) Western blot analysis of global lactylation levels in OFs transfected with the indicated siRNAs and stimulated with or without M22 (n = 3). β-actin served as the loading control. (I) Predicted binding site of CREB at the LDHA promoter by JASPAR online tools. (J) Analysis of the binding affinity between CREB and the LDHA promoter. The schematic diagram of the EMSA probe design (left panel), and the resulting EMSA gel (right panel). (K) High-content imaging analysis of LDHA-tdTomato expression in OFs treated with or without M22. Representative images are shown in the left panel (scale bars = 100 μm). The right panel displays violin plots quantifying tdTomato fluorescence intensity as determined by the high-content imaging system. (L) The mRNA expression levels of LDHA in OFs transfected with the indicated siRNAs and stimulated with or without M22 (n = 3). (M) Western blot analysis of CREB and LDHA in OFs transfected with indicated siRNAs (n = 3). β-actin served as the loading control. (N) Lactate levels in OFs transfected with the indicated siRNAs and stimulated with or without M22 (n = 3). (O) Western blot analysis of the global lactylation levels of OFs transfected with indicated siRNAs followed by 48 h of stimulation with M22 (n = 3). β-actin served as the loading control. All data are represented as the mean ± SEM. Two-tailed unpaired Student’s t-tests were performed in (B, C, D, F, K). One-way ANOVA, followed by Tukey’s multiple post hoc test, was performed in (E, G, H, L-O). Accurate *p*-values are listed in the figures.

The primary target of TSAb is the thyrotropin receptor (TSHR), a G protein-coupled receptor (GPCR). TSHR activation triggers the cAMP-PKA-CREB signaling axis, promoting proinflammatory cytokines and hyaluronan production, thereby perpetuating tissue inflammation and edema^18–20^. Consistent with TSHR-mediated activation of this axis, M22 treatment elevated intracellular cAMP levels and increased the p-CREB/CREB ratio in OFs (Supplementary Fig. 6A-B). TSHR also engages in crosstalk with other receptors, such as the insulin-like growth factor-1 receptor (IGF-1R) in TED, thereby amplifying systemic inflammatory responses and metabolic signaling^4, 21^. CREB is a nuclear transcription factor downstream effector of the intracellular cAMP/PKA signaling that binds cAMP response elements (CRE) in target promoters to regulate gene transcription^22^. Given that M22 increases LDHA mRNA levels, we hypothesized that CREB directly promotes LDHA transcription. Using the JASPAR database, we identified two putative CRE binding sites within the LDHA promoter region (Fig. 3I). An electrophoretic mobility shift assay (EMSA) confirmed that CREB binds to the LDHA probe, producing a retarded migration band (Fig. 3J). To test whether M22 promotes LDHA transcription, we constructed a tdTomato reporter plasmid. High-content imaging showed higher integrated fluorescence intensity in M22-treated cells than in controls (Fig. 3K). To examine whether CREB regulates LDHA, we used siRNA to knock down CREB expression. All three si-CREB sequences effectively reduced CREB at both the mRNA and protein levels (Supplementary Fig. 5C-D). Consistently, CREB silencing significantly downregulated LDHA expression at both the mRNA and protein levels (Fig. 3L-M). This decrease was accompanied by reduced intracellular lactate content and global lactylation levels (Fig. 3N-O). Together, these findings indicate that TSAbs drive lactate production and protein lactylation through CREB-mediated transcriptional activation of LDHA.

### 2.4. TSAbs promote adipogenesis via a lactate-mediated mechanism

To determine whether lactate mediates the pro-adipogenic effects of M22, we treated GOFs with oxamate. Oxamate abolished M22-induced lipid droplet accumulation, as assessed by Oil Red O staining, and reduced BODIPY-stained neutral lipids (Supplementary Fig. 7A-C). Oxamate also attenuated M22-induced triglyceride production and blocked the upregulation of PPARγ and C/EBPα protein levels (Supplementary Fig. 7D-F). To rule out off-target effects of pharmacological inhibition, we knocked down LDHA using siRNA. Consistently, LDHA silencing reversed M22-induced lipid droplet accumulation and blunted the increase in neutral lipid content (Fig. 4A-C), suppressed triglyceride production, and prevented the upregulation of PPARγ and C/EBPα protein levels (Fig. 4D-E).

**Figure 4.**
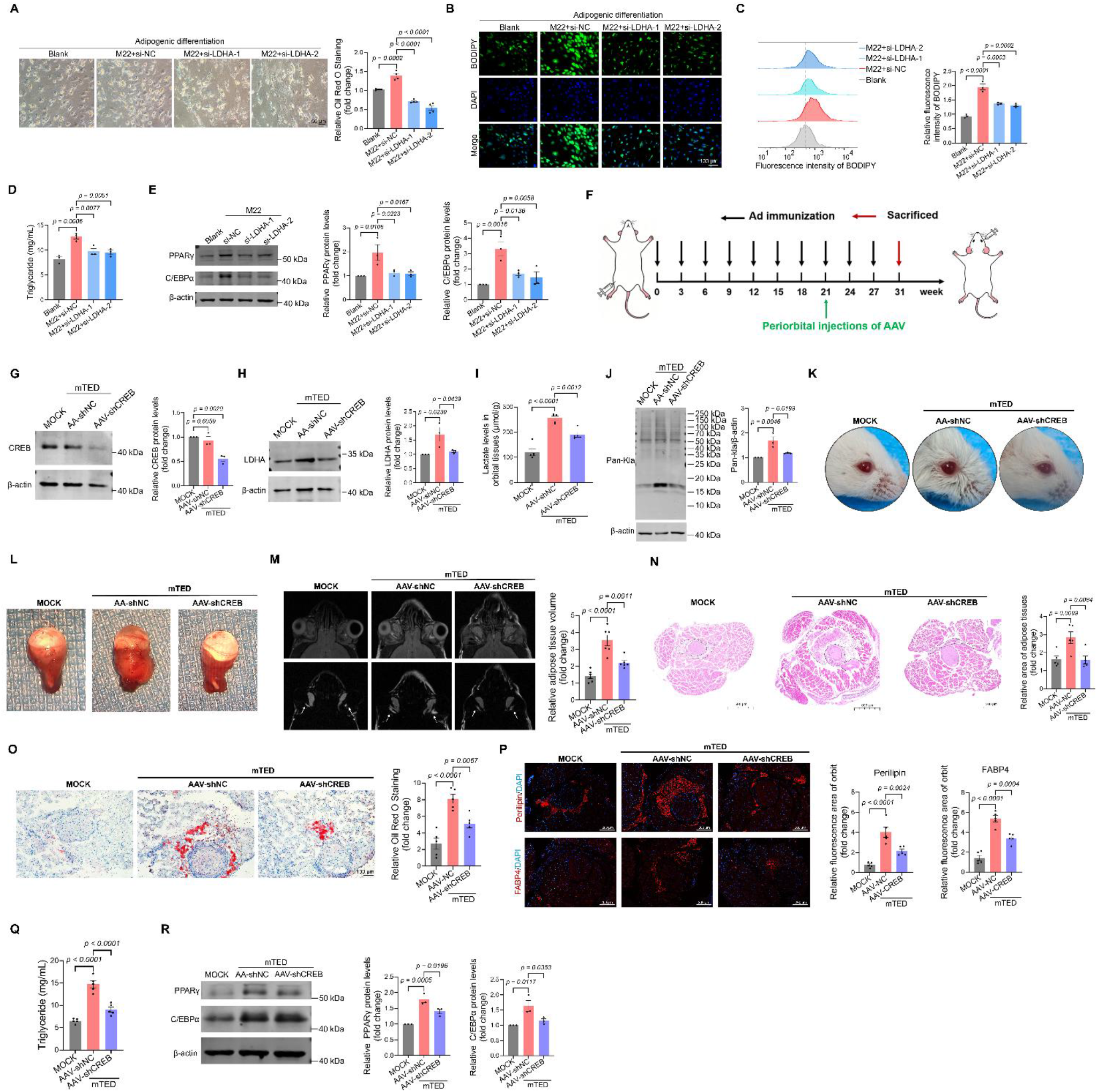
TSAbs promote adipogenesis via a lactate-mediated mechanism. (A) Representative Oil Red O staining images of adipogenic-differentiated OFs transfected with the indicated siRNAs and stimulated with or without M22. Scale bar = 50 μm. (B) Representative BODIPY staining images of adipogenic-differentiated OFs transfected with the indicated siRNAs and stimulated with or without M22. Scale bar = 100 μm. (C) Flow cytometry analysis of adipogenic-differentiated OFs transfected with the indicated siRNAs and stimulated with or without M22 (n = 3). (D) Triglyceride content in adipogenic-differentiated OFs transfected with the indicated siRNAs and stimulated with or without M22 (n = 3). (E) Western blot analysis of PPARγ and C/EBPα in adipogenic-differentiated OFs transfected with the indicated siRNAs and stimulated with or without M22 (n = 3). β-actin served as the loading control. (F) Experimental timeline showing the schedule for model construction and periorbital AAV administration in mice. (G) Western blot analysis of CREB protein levels in orbital tissues from the three mouse groups (n = 3). β-actin served as the loading control. (H) Western blot analysis of LDHA protein levels in orbital tissues from the three mouse groups (n = 3). β-actin served as the loading control. (I) Lactate levels were measured in the orbital tissues of mice from the three mouse groups (n = 5). (J) Western blot analysis of the global lactylation levels of mice orbital tissues from the three mouse groups (n = 3). (K) Representative images of the mice’s ocular surface across the three mouse groups. (L) Representative images of mice’s orbits across the three mouse groups. (M) Representative images showing T2-weighted magnetic resonance imaging (upper panels) and fat-water separation maps (lower panels) for mice in the three mouse groups. Corresponding quantitative analysis of relative adipose tissue volume based on MRI (n = 6). (N) Representative hematoxylin-eosin (H&E) staining images of orbital tissue sections from the four mouse groups and the corresponding quantitative analysis of the relative area of adipose tissues (n = 5). ON, optic nerve. The region outlined by the black dashed line represents adipose tissue. Scale bar=500 μm. (O) Representative Oil Red O staining images of orbital tissue sections from the three mouse groups (n = 5). Scale bar = 100 μm. (P) Representative immunofluorescence images of Perilipin and FABP4 in orbital tissue sections from the three mouse groups and corresponding quantitative analysis of fluorescence area (n = 5). Scale bar = 100 μm. (Q) Triglyceride content in the orbits from the three mouse groups. (R) Western blot analysis of PPARγ and C/EBPα protein levels in orbital tissues from the three mouse groups (n = 3). β-actin served as the loading control. All data are represented as the mean ± SEM. Two-tailed unpaired Student’s t-tests were performed in (F). One-way ANOVA, followed by Tukey’s multiple post hoc test, was performed in (A, C-E, G-J, M-R). Accurate *p*-values are listed in the figures.

We further extended these findings in vivo using AAV-mediated shRNA delivery to reduce LDHA expression in mouse orbits. Given that the transcription factor CREB positively regulates LDHA expression, we delivered periorbital injections of AAV-shCREB targeting CREB. The experimental timeline is shown in Fig. 4F. After nine adenoviral injections to establish immune-induced TED, mice were randomly assigned to receive AAV-shNC (mTED+AAV-shNC) or AAV-shCREB (mTED+AAV-shCREB). Among the three tested shRNA sequences, AAV-shCREB-2 showed the highest knockdown efficiency and was selected for subsequent experiments (Supplementary Fig. 8). Western blotting confirmed that AAV-shCREB administration reduced both CREB and LDHA protein levels in mouse orbital tissues (Fig. 4G-H), leading to a significant decrease in lactate content and pan-lactylation levels (Fig. 4I-J). Consistent with these molecular changes, AAV-shCREB-treated mice showed reduced eyelid swelling and congestion (Fig. 4K). Orbital imaging further demonstrated that AAV-shCREB reduced soft tissue expansion (Fig. 4L). T2-weighted MRI with water-fat separation quantified adipose tissue volume, which was increased in the mTED+AAV-shNC group and markedly reduced following AAV-shCREB administration (Fig. 4M). Histological and molecular analyses confirmed these observations. H&E and Oil Red O staining showed that adipose tissue area was expanded in the mTED+AAV-shNC group but markedly reduced in the AAV-shCREB group (Fig. 4N-O). Immunofluorescence staining for the adipogenic markers Perilipin and FABP4 confirmed this reduction in adipose tissue (Fig. 4P). Consistent with these findings, biochemical assays revealed that triglyceride levels were elevated in the mTED+AAV-shNC group but significantly reduced after AAV-shCREB treatment (Fig. 4Q). At the molecular level, PPARγ and C/EBPα expression was upregulated in the mTED+AAV-shNC group and downregulated in the mTED+AAV-shCREB group (Fig. 4R). Together, these results indicate that TSAbs promote adipogenesis through a lactate-dependent mechanism.

### 2.5. H3K18la is upregulated by TSAb stimulation and promotes adipogenesis

We previously observed that global lactylation was elevated in GOFs and that lactylated proteins were enriched at approximately 15 kDa and localized to the nucleus (Fig. 1L, 5A). Given that core histones have molecular masses of ∼11–15 kDa and are the most abundant nuclear proteins, this pattern raised the possibility that histone lactylation may contribute to the observed signal. To test this hypothesis, we quantified lactylation levels at seven known histone sites (H3K9la, H3K14la, H3K18la, H4K5la, H4K8la, H4K12la, and H4K16la) in OFs following M22 treatment. Among these, H3K18la levels showed the most pronounced increase (Fig. 5B-C). Confocal imaging confirmed that H3K18la fluorescence intensity was increased upon M22 treatment (Fig. 5D), and co-staining revealed strong colocalization between lactylation and histone H3, with Pearson’s correlation coefficients and overlap coefficients each exceeding 0.8 (Fig. 5E). To assess H3K18la levels in diseased tissues, we performed western blotting and IHC on orbital tissues from TED patients and TED mice. H3K18la levels were significantly elevated in both TED patients and TED mice compared with normal controls (Fig. 5F-I). To determine whether H3K18la is functionally required for adipogenesis, we generated a histone H3K18R mutant that cannot be lactylated. The plasmid schematic is shown in Fig. 5J. Immunoprecipitation assays confirmed that expression of the H3K18R mutant prevented the M22-induced increase in H3K18la (Fig. 5K). Functionally, OFs expressing the H3K18R mutant displayed impaired lipid accumulation and reduced triglyceride content compared with those expressing wild-type H3 (Fig. 5L-M). Consistent with this defect, H3K18R mutant expression led to downregulation of PPARγ and C/EBPα protein levels (Fig. 5N). These results indicate that H3K18la is a downstream effector of lactate-mediated adipogenesis in TED.

**Figure 5.**
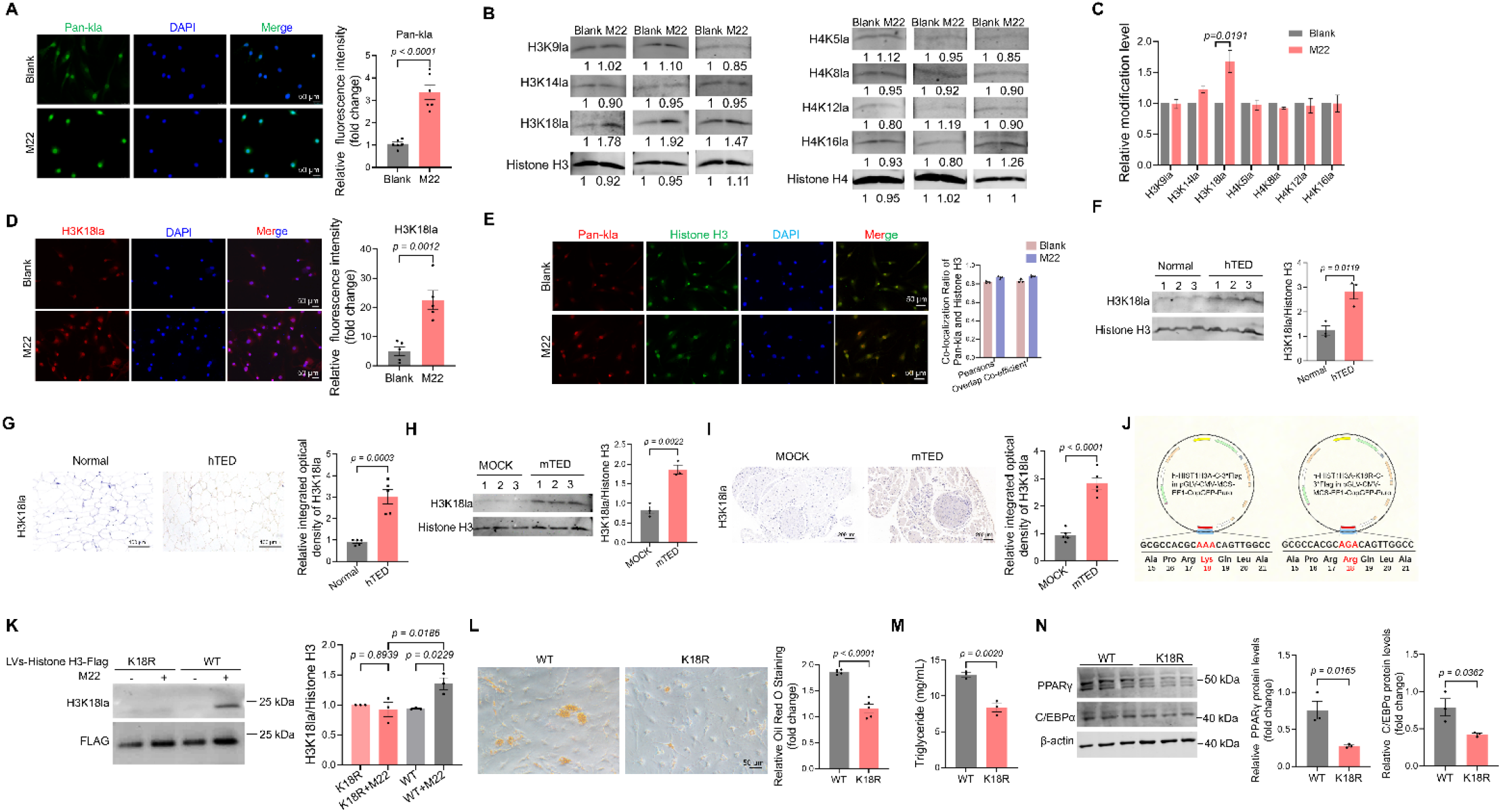
H3K18la is upregulated by TSAb stimulation and promotes adipogenesis. (A) Representative immunofluorescence images of Pan-kla (green) in OFs (left panel, scale bar = 50 μm) and corresponding quantitative analysis of relative fluorescence intensity (right panel, n = 6). (B) Western blot analysis of histone lactylation in OFs with or without M22 treatment (n = 3). Histone H3 and Histone H4 served as the loading controls. (C) The corresponding quantitative analysis of the relative histone lactylation level of panel (B). (D) Representative immunofluorescence images of H3K18 lactylation (red) in OFs stimulated with or without M22 (left panel, scale bar = 50 μm) and corresponding quantitative analysis of relative fluorescence intensity of H3K18la (right panel, n = 5). (E) Representative confocal images of Pan-kla (red) and Histone H3 (green) in OFs (left panel, scale bar = 50 μm). Colocalization analysis using Pearson’s correlation coefficient and Overlap coefficient of Pan-kla and Histone H3 in OFs (right panel, n = 3). (F) Western blot analysis of H3K18 lactylation in orbital tissues from normal donors (n = 3) and TED patients (n = 3). (G) IHC staining images of H3K18 lactylation in orbital adipose tissues from normal donors (n = 5) and TED patients (n = 5) and corresponding quantitative analysis of relative integrated optical density. Scale bar = 100 μm. (H) Western blot analysis of H3K18 lactylation in orbital tissues from MOCK and mTED groups (n = 3). (I) IHC staining images of Pan-kla in mouse orbital tissues from the MOCK and mTED groups and corresponding quantitative analysis of relative integrated optical density (n = 5). Scale bar = 100 μm. (J) Plasmid maps of HIST1H3A-WT and HIST1H3A-K18R expression vectors. (K) Western blot analysis of PPARγ and C/EBPα in OFs expressing HIST1H3A-WT or HIST1H3A-K18R following adipogenic differentiation and Nala treatment (n = 3). (L) Representative Oil Red O staining of OFs expressing HIST1H3A-WT or HIST1H3A-K18R following adipogenic differentiation and Nala treatment (n = 5). Scale bar = 50 μm. (M) Triglyceride content in OFs expressing HIST1H3A-WT or HIST1H3A-K18R following adipogenic differentiation and Nala treatment (n = 3). (N) Western blot analysis of PPARγ and C/EBPα protein levels OFs expressing HIST1H3A-WT or HIST1H3A-K18R following adipogenic differentiation and Nala treatment (n = 3). β-actin served as the loading control. All data are represented as the mean ± SEM. Two-tailed unpaired Student’s t-tests were performed in (A, C, D, F-H, L-N). One-way ANOVA, followed by Tukey’s multiple post hoc test, was performed in (K). Accurate *p*-values are listed in the figures.

### 2.6. H3K18la promotes adipogenesis by transcriptionally activating PPARγ and C/EBPα

Histone modifications play an important role in regulating the transcription of target genes^23^. To identify genes regulated by H3K18la in OFs, we performed CUT&Tag analysis using an anti-H3K18la antibody (Fig. 6A). H3K18la was enriched in the promoter and upstream regions, with higher enrichment in the M22 group than in the control group (Fig. 6B). Over 20% of the total H3K18la binding peaks were localized within ±1 kb of the transcription starts sites (TSS) (Supplementary Fig. 9A-B). We identified 1561 differentially enriched peaks between the M22-treated and control groups. Pearson’s correlation analysis confirmed high reproducibility among biological replicates (Supplementary Fig. 9C), and peaks were evenly distributed across all chromosomes (Supplementary Fig. 9D). Pathway enrichment analysis of genes associated with differentially bound peaks (fold change < 0.67 or > 1.5) revealed that genes with upregulated H3K18la peaks were enriched in lipid metabolism-related pathways, including adipogenesis, fatty acid metabolic process regulation, fat cell differentiation, lipid biosynthetic process, and lipid and atherosclerosis (Fig. 6C). CUT&Tag showed increased H3K18la at the promoters of PPARG and CEBPA in M22-stimulated OFs (Fig. 6D), and CUT&Tag-qPCR analysis confirmed that M22-induced hyperlactylation increased H3K18la enrichment at these promoters (Fig. 6E). Consistently, both mRNA and protein levels of PPARγ and C/EBPα were upregulated in M22-stimulated OFs (Fig. 6F-G). Together, these results indicate that H3K18la promotes adipogenesis by transcriptionally activating PPARγ and C/EBPα.

**Figure 6.**
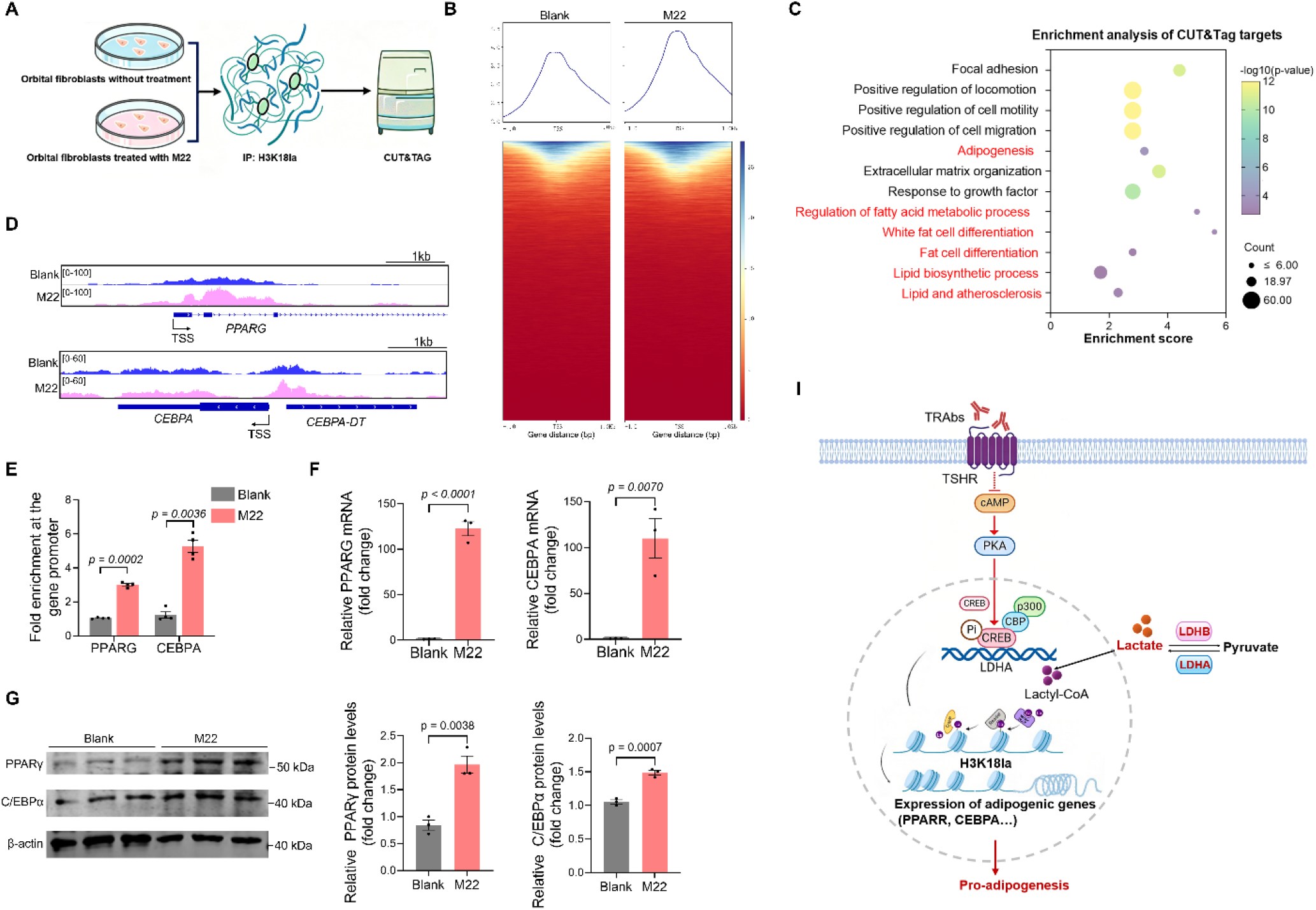
H3K18la promotes adipogenesis by transcriptionally activating PPARγ and C/EBPα. (A) Schematic representation of CUT&Tag analysis. (B) The binding density of H3K18la in OFs, with or without M22 treatment, was visualized using deepTools, and the read distribution relative to the TSS is presented as a line chart. (C) Enrichment analysis of elevated H3K18la binding peaks at candidate target genes. (D) Genome browser tracks of CUT&Tag signals at the PPARG and CEBPA loci, visualized using IGV. (E) Fold enrichment of H3K18la at the PPARG and CEBPA gene promoters analyzed by CUT&Tag-qPCR. (F) qRT-PCR assays monitoring expression of PPARG and CEBPA in OFs treated with or without M22. (G) Expression of PPARγ and C/EBPα in M22-treated and control OFs was determined by western blotting. (I) Schematic summary of adipogenesis induced by the TSHR/CREB/LDHA axis-mediated histone lactylation. All data are represented as the mean ± SEM. Two-tailed unpaired Student’s t-tests were performed in (E-G). Accurate *p*-values are listed in the figures.

## 3. Discussion

The intersection of immune dysregulation and metabolic perturbation is increasingly recognized as a driving force in autoimmune pathogenesis, with the Warburg effect, a metabolic shift toward aerobic glycolysis, at its core^24, 25^. In TED, TSAbs engage the TSHR on OFs to initiate disease, yet how this immune trigger is coupled to metabolic reprogramming and subsequent tissue remodeling remains incompletely understood. Here, we demonstrate that lactate, the terminal glycolytic metabolite, serves as a central hub in TSHR-mediated adipogenesis, functioning through a CREB/LDHA-dependent cascade that culminates in H3K18 lactylation and transcriptional activation of adipogenic programs.

TED has traditionally been viewed as an organ-specific autoimmune disorder driven by autoreactive lymphocytes and aberrant activation of orbital fibroblasts. Although this immune-centric model is supported by extensive evidence, including the identification of TSHR and IGF-1R as key autoantigens^26^, it does not fully explain how immune activation translates into the adipogenic and tissue remodeling phenotypes that define the disease. Our findings reveal that metabolic reprogramming, specifically the TSHR-CREB-LDHA-lactate cascade and subsequent H3K18 lactylation, serves as a critical intermediary that couples autoimmune signaling with adipogenesis. In this framework, TSAb stimulation drives a shift toward aerobic glycolysis in orbital fibroblasts, generating lactate that fuels histone lactylation and the transcriptional activation of adipogenic programs. Rather than being a mere downstream consequence of inflammation, metabolic reprogramming appears to actively participate in disease progression. By framing TED as an immune-metabolic disorder, this integrative perspective reconciles the centrality of autoimmune mechanisms with the metabolic alterations that shape the disease phenotype, potentially offering a more comprehensive framework for identifying therapeutic targets.

TSHR is a G protein-coupled receptor (GPCR) that orchestrates diverse signaling pathways involved in metabolic regulation in multiple tissues^27, 28^. Our study identified the transcription factor CREB as a molecular node that bridges TSHR signaling and metabolic reprogramming. While CREB is well-established as a downstream effector of the TSHR-cAMP-PKA axis and has been implicated in the transcriptional regulation of inflammatory mediators in TED^20^, we demonstrate that CREB also directly activates LDHA transcription, coupling receptor engagement to lactate production. This positions CREB not only as a regulator of inflammation but also as a molecular link between early immune activation and the metabolic reprogramming that drives tissue remodeling.

The relationship between lactate and lipid metabolism is highly tissue-specific and operates through dual mechanisms: lactate serves as both a metabolic substrate for triglyceride synthesis and a signaling molecule that inhibits lipolysis via GPR81-mediated suppression of cAMP/PKA signaling^29, 30^. Our findings reveal a third mechanism by which lactate promotes adipogenesis in the orbit via histone lactylation. Lactate functions as an epigenetic modulator rather than acting solely as a metabolic substrate. This lactylation-dependent mechanism differs fundamentally from the substrate-driven and receptor-mediated pathways described previously, expanding our understanding of how lactate regulates lipid metabolism at the chromatin level.

Several limitations of this study should be considered. First, although our functional analyses focused on adipogenesis, the contribution of lactate and H3K18la to other pathological features of TED, including inflammation, tissue edema, and extraocular muscle involvement, requires further study. Second, the H3K18R mutation, although it prevents lactylation at this site, could potentially alter nucleosome structure or crosstalk with modifications at neighboring residues. Finally, validation in larger and more diverse cohorts will be necessary to establish the generalizability of lactate and H3K18la as biomarkers.

In summary, this study reveals that in TED, TSAbs trigger a TSHR-CREB-LDHA-lactate-H3K18la axis, where lactate accumulation drives H3K18 lactylation and subsequent adipogenesis. These findings highlight metabolic reprogramming via immune-epigenetic crosstalk as a core pathogenic mechanism, establishing TED as an immune-metabolic disorder and identifying the lactate-H3K18la axis as a promising therapeutic target.

## 4. Experimental Section

### Study approval

This study was approved by the Ethics Committee of Wenzhou Medical University and adhered to the principles of the Declaration of Helsinki. Informed consent was obtained from all the participants. Human tissue specimens were collected with ethical approval (No. 2023-074-K-62). Human tissue experiments complied with the guidelines of the ARVO Best Practices for Using Human Eye Tissue in Research (Nov2021). Animal experiments were approved by the Institutional Animal Care and Use Committee (No. YSG226041302) and conducted in accordance with the institutional guidelines and the *Guide for the Care and Use of Laboratory Animals* guidelines.

### Patients and samples

OATs were obtained from eighteen patients with TED undergoing orbital decompression surgery for severe disease, as well as from thirteen healthy donors undergoing blepharoplasty. All procedures were performed at the Eye Hospital of Wenzhou Medical University (Wenzhou, China). The diagnosis of TED was established according to the 1995 Bartley criteria^31^. Disease activity and severity were evaluated using the Clinical Activity Score (CAS)^32^ and guidelines from the European Group on Graves′ Orbitopathy (EUGOGO)^33^. All TED patients presented with moderate-to-severe disease activity (CAS < 4) and had discontinued steroid or immunosuppressive therapy for at least three months prior to surgery. None of the patients had a history of radioactive iodine therapy or other systemic or ocular autoimmune conditions. Healthy donors had no history of autoimmune or inflammatory diseases, thyroid or orbital disorders, or prior orbital surgeries. Written informed consent was obtained from all participants.

### Cells culture

OFs were isolated from the orbital adipose tissues of both TED patients and healthy donors by mincing the tissue into small fragments, which were then plated onto culture dishes. After allowing time for cell attachment, the explants were cultured in high-glucose Dulbecco’s Modified Eagle’s Medium (DMEM, Gibco, USA) supplemented with 20% fetal bovine serum (FBS) and 1% penicillin/streptomycin (P/S), maintained at 37°C in a 5% CO₂ atmosphere. All functional assays were performed using OFs at passages 3-7, and each experiment was independently repeated with at least three distinct donor-derived cell lines.

### Adipogenesis

When the OFs were confluent to 100%, the complete growth medium was converted to a differential medium (DM). The base medium for DM was high-glucose DMEM containing 10% FBS and 1% P/S, supplemented with 0.1 mM indomethacin (Sigma-Aldrich, Germany), 0.1 μM dexamethasone (Sigma-Aldrich, Germany), 0.5 mM 3-isobutyl-1-methylxanthine (IBMX) (MedChemExpress, China), and 10 μg/mL insulin (Procell, China). The medium was refreshed every 4-5 days, and the induction period lasted for 10-14 days, as previously described^34^.

### siRNA transfection assay

Cells were seeded into 6-well plates at a density of 2 × 10^5^ cells per well and cultured for 24 h. siRNA was diluted in 250 µL of Opti-MEM medium (11058021, Gibco, USA) and mixed with Lipofectamine™ 3000 (L3000015, Thermo Fisher Scientific, USA), which had been separately diluted in 250 µL of Opti-MEM medium. The resulting transfection complex was then added dropwise to each well. After 6 h of incubation, the transfection mixture was removed and replaced with fresh basal medium, and cells were cultured for an additional 48 h. Subsequently, cells were harvested for western blotting and RT-qPCR analyses.

### Single-cell RNA-seq and bioinformatics analysis

To analyze the differential expression of glycolysis-related genes, raw single-cell RNA sequencing (scRNA-seq) data from normal and TED orbital tissues were retrieved from the Gene Expression Omnibus (GEO) database under accession number GSE194323^35^. Glycolysis/gluconeogenesis-specific KEGG pathway annotation (hsa00010) was employed as a framework to construct heatmaps. Data normalization was performed using the limma package (v3.50.0) with Relative Log Expression (RLE) transformation. Glycolysis-related gene expression was visualized using z-score-scaled heatmaps generated with the heatmap (v1.0.12). All computational analyses were conducted in R version 4.3.1.

### Expression vector construction and transfection

The human HIST1H3A-C-3Flag expression construct (NM_003529.3; GeneCopoeia) was cloned into a pcDNA3.1-P2A-EGFP vector to generate a wild-type (WT) plasmid. The K18R mutant construct (HIST1H3A-C-3Flag K18R) was cloned into the same vector. All constructs were verified using DNA sequencing. Both plasmids were co-transfected with lentiviral packaging vectors to produce recombinant particles. OFs were subsequently infected with these viral particles and subjected to puromycin selection to establish stable HIST1H3A-WT and HIST1H3A-K18R overexpressing OFs cell lines. All plasmid constructions and lentiviral packaging were performed by Ji’an Biotechnology Co., Ltd.

### Metabolomics

80 mg of each sample was mixed with isotope internal standards and cold methanol/acetonitrile solution (1:1, v/v). The lysate was homogenized by MP homogenizer, adequately vortex, then ultrasounded for 5 min at low temperature, followed by incubation at -20℃ for 1 h. The mixture was centrifuged for 20 min. The supernatant was taken over the Ostro plate, and the receiving solution was divided into two tubes (one tube is 2/3 and the other tube is 1/3). 2/3 of the receiving solution was dried in a vacuum centrifuge, the samples were re-dissolved in acetonitrile/water (1:1, v/v) and adequately vortexed, and then centrifuged. The supernatants were collected for LC-MS/MS analysis. Analyses were performed using an UHPLC (LC-30AD, Shimadzu) coupled to a QTRAP (AB Sciex 6500+). Chromatographic separation was performed on both Amide (35°C) and C18 (40°C) columns using specific gradient elution profiles with ammonium acetate/ammonia and formic acid modifiers, respectively. Quality control samples and standard metabolite mixtures were utilized to monitor system stability and correct retention times, while Multiquant software was employed for peak extraction and data processing by Hangzhou Astrocyte Technology Co., Ltd.

### Histopathological examination

For H&E staining, the sections were stained with hematoxylin, rinsed, differentiated, and rinsed again. The slides were then immersed in ammonia solution, rinsed, dehydrated with 85% and 95% ethanol, and stained with eosin. After dehydration with a gradient of alcohol, the slides were cleared with xylene and sealed with neutral gum. Image data were collected using a Pannoramic Slide Scanner (3DHISTECH, Budapest, Hungary) and a DM4B biological microscope (Leica, Bannockburn, IL, USA). The average area of the adipose tissue surrounding the optic nerve was quantified using Fiji/Image J software. The adipose tissue area of each slice was standardized with respect to the optic nerve area.

### Immunohistochemistry

Tissue sections were deparaffinized in xylene and rehydrated using a graded ethanol series. Antigen retrieval was conducted by incubating the sections in citrate buffer (pH 6.0), followed by quenching of endogenous peroxidase activity at room temperature. The sections were blocked with 3% bovine serum albumin (BSA) for 1 hour, and then incubated with the appropriate primary antibodies overnight at 4°C. Subsequently, the samples were incubated with horseradish peroxidase (HRP)-conjugated secondary antibodies for 1 hour at room temperature. Immunoreactivity was visualized using 3,3’-diaminobenzidine tetrahydrochloride (DAB) as the chromogen, followed by a 3-minute hematoxylin counterstain. After dehydration through a graded ethanol series and clearing in xylene, the sections were mounted with neutral balsam. Whole-slide images were acquired using a Pannoramic Slide Scanner (3DHISTECH, Budapest, Hungary) and a DM4B biological microscope (Leica, Bannockburn, IL, USA). The integrated optical density (IOD) of the stained regions was quantified using Fiji/Image J software.

### Immunofluorescence

Cells were fixed with 4% paraformaldehyde, washed, and subsequently blocked using 3% BSA. The samples were incubated with primary antibodies overnight at 4°C. Following extensive washing, cells were incubated with the appropriate fluorescent-conjugated secondary antibodies and counterstained with DAPI to visualize the nuclei. Fluorescence images were acquired using a Cell Observer SD system (Zeiss, Germany).

Tissue sections were deparaffinized in xylene and rehydrated through a graded series of ethanol solutions. Antigen retrieval was performed by incubating the sections in citrate buffer (pH 6.0 and blocking with 3% BSA. The sections were then incubated with primary antibodies overnight at 4°C. After washing, fluorescent-conjugated secondary antibodies were incubated, followed by DAPI nuclear counterstaining. Images were acquired using a DM4B biological microscope (Leica, Bannockburn, IL, USA).

### Lactate measurement

The concentrations of lactate in orbital tissues and OFs were detected using an L-lactate assay kit (KTB1100, Abbkine, China) according to the manufacturer’s instructions. Quantification was performed using a microplate reader (Molecular Devices) at 450 nm.

### Oil Red O staining

Oil Red O staining of OFs was performed using an ORO kit (PHYGENE, China) according to the manufacturer’s protocol. Stained cells were visualized under an inverted microscope (Nikon, Japan). For quantification, the dye was solubilized with isopropanol, and the optical density (OD) was measured at 490 nm using a microplate reader (Molecular Devices). For orbital tissue sections, the slides were incubated with Oil Red O solution (O0625, Sigma-Aldrich, Germany) for 30 min at room temperature in the dark. Excess dye was removed by rinsing with 60% isopropanol, followed by hematoxylin counterstaining. High-resolution images were acquired using a Pannoramic Slide Scanner (3DHISTECH, Budapest, Hungary), and the ORO-positive area was quantified using Fiji/ImageJ software.

### BODIPY staining

Cells were seeded into 6-well plates and cultured until they reached 60–80% confluence. After rinsing twice with phosphate-buffered saline (PBS, pH 7.4), cells were incubated with BODIPY 493/503 (5μM) (C2053S, Beyotime, China) for 30 min at 37°C in a 5% CO_2_ incubator. All steps were performed under light protection to avoid fluorescence quenching. Following staining, cells were gently washed 2–3 times with PBS to remove unbound dye, then counterstained with Hoechst 33342 (1 µg/mL in PBS) for 5 min at room temperature to label cell nuclei. Fluorescent images were acquired immediately using a DMi8 biological microscope (Leica, Bannockburn, IL, USA). For quantitative analysis, cells were harvested, and fluorescence signals were detected using an Attune NxT flow cytometer (Thermo Fisher Scientific, USA). Flow cytometric data were analyzed using FlowJo 10.0 software. All experiments were independently repeated thrice.

### Triglyceride assay

Triglyceride levels were quantified using the Amplex Red Triglyceride Assay Kit (S0219S, Beyotime, China). Lipids were extracted from cells and tissue samples via isopropanol homogenization. The extracts were then diluted 5-fold according to the manufacturer’s protocol and incubated with the triglyceride detection working solution in the dark. Fluorescence intensity was measured using a fluorescence microplate reader with excitation and emission wavelengths set at 530 nm and 590 nm, respectively.

### RNA isolation, reverse transcription and quantitative RT-PCR

Total RNA was isolated from cultured cells and orbital adipose tissues using TRIzol reagent (15596026, Invitrogen, USA) in accordance with the manufacturer’s protocol. RNA concentration and purity were assessed using a UV spectrophotometer. Complementary DNA (cDNA) was synthesized with a reverse transcription kit (Promega, USA), and quantitative real-time polymerase chain reaction (qRT-PCR) was performed using HotStart™ Universal 2×Green qPCR Master Mix (K1170, APExBIO, USA). All experiments were independently repeated three times.

### Electrophoretic mobility shift assay

Electrophoretic mobility shift assay (EMSA) was performed using a Chemiluminescent EMSA Kit (GS009, Beyotime, China) strictly following the manufacturer’s instructions. HPLC-purified, biotin-labeled double-stranded DNA probes targeting the LDHA promoter, along with unlabeled specific and nonspecific competitor probes, were synthesized by Sangon Biotech (Shanghai, China) and annealed prior to use. Recombinant human CREB1 protein (P0471, FineTest, China; 2 μg per reaction) was incubated with the biotin-labeled probe in a 20 μL binding reaction containing 1X binding buffer. Parallel reactions included a negative control (no protein) and a specificity control (100-fold molar excess of unlabeled specific probe). Following a 30-minute incubation at room temperature in the dark, 5 μL of 5X EMSA/Gel-Shift loading buffer (blue) was added to each reaction. The samples were loaded onto a pre-run 4% non-denaturing polyacrylamide gel in 0.5X TBE, alongside 10 μL of diluted 1X loading buffer in an adjacent lane to monitor migration. Electrophoresis was conducted at 120 V at 4°C until the bromophenol blue dye front migrated approximately two-thirds of the gel length. Subsequently, the DNA-protein complexes were transferred to a positively charged nylon membrane via electroblotting and cross-linked using a UV crosslinker. Finally, biotin-labeled complexes were detected using a chemiluminescent substrate and visualized with an imaging system.

### Seahorse assay

The oxygen consumption rate (OCR) of OFs was assessed using a Seahorse XF96 Pro Analyzer (Agilent Technologies, USA) following the manufacturer’s standard protocol. Cells were seeded into XF96 microplates and cultured to approximately 90% confluence. Prior to the assay, the growth medium was replaced with Seahorse assay medium, and the plates were incubated at 37°C in a CO₂-free incubator for 1 hour to equilibrate. Glycolytic function was evaluated using the Seahorse XF Glycolysis Stress Test Kit (103015-100, Agilent Technologies, USA) through the sequential injection of glucose (10 mM), oligomycin (1 μM), and 2-deoxy-D-glucose (50 mM). Data acquisition and analysis were performed using Wave software (Agilent Technologies), and all resulting parameters were normalized to the cell count per well at the time of measurement.

### CUT&Tag assay and CUT&Tag-qPCR

CUT&Tag assays were performed using a commercial kit (HD102-01, Vazyme) according to the manufacturer’s instructions. OFs treated with or without M22 were immobilized on Concanavalin A-coated magnetic beads, washed, and incubated overnight at 4°C with rotation using anti-H3K18la antibody (PTM1406RM, PTMBio, China). After washing, cells were incubated with a secondary antibody for 1 hour at room temperature to tether the protein A– Tn5 (pA-Tn5) fusion enzyme. Following the addition of pA-Tn5 and a 1-hour incubation, tagmentation was initiated by adding magnesium-containing buffer. The reaction was stopped using the provided stop buffer. DNA was purified using the kit’s buffer, amplified via PCR, and libraries were purified (VAHTS DNA Clean Beads) and quantified (VAHTS Library Quantification Kit) prior to paired-end sequencing on an Illumina Novaseq platform.

CUT&Tag-qPCR was performed using a commercial kit (TD904-C1, Vazyme) according to the manufacturer’s instructions. After extraction, amplification, and purification, the DNA was subjected to sequencing and qPCR. The resulting data were analyzed using the 2^-△△CT^ method to determine the relative fold changes.

### Western blotting

OFs or orbital adipose tissues were lysed in RIPA lysis buffer with a protease inhibitor cocktail (EpiZyme, China) and acetylase inhibitor cocktail (Beyotime, China). Protein concentrations were determined using a BCA Protein Assay Kit (EpiZyme, China). Equal amounts of protein samples were separated via sodium dodecyl sulfate-polyacrylamide gel electrophoresis (SDS-PAGE) and subsequently transferred onto nitrocellulose membranes using a Bio-Rad Western blot transfer system. After blocking, the membranes were incubated with the corresponding primary antibodies at 4°C overnight, followed by incubation with fluorescence-conjugated secondary antibodies (1:5000 dilution, LI-COR, USA) at room temperature for 1 h. Protein bands were visualized and imaged using an Odyssey infrared scanning system (LI-COR, USA). The gray values of the target protein bands were quantified using the ImageJ software.

### Immunoprecipitation assay

OFs were lysed using a lysis buffer supplemented with protease inhibitors (P1051, Beyotime, China). The resulting lysates were clarified by centrifugation and pre-cleared by incubation with control IgG-conjugated Protein A/G magnetic beads at 4°C for 1 h to minimize non-specific binding. The pre-cleared supernatants were incubated overnight at 4°C with gentle rotation in the presence of Protein A/G magnetic beads conjugated with an anti-DYKDDDDK tag primary antibody (66008-4-Ig, Proteintech, China). Immunoprecipitated complexes were captured magnetically, washed multiple times with lysis buffer to remove non-specifically bound proteins, and eluted by boiling in SDS loading buffer for downstream western blotting analysis.

### Establishment of TED mouse model

The TED mouse model was established as described previously^16^. Female BALB/c mice were acquired from Weitong Lihua Experimental Animal Technology Co., Ltd. and accommodated in a specific pathogen-free facility at the Eye Hospital of Wenzhou Medical University. Mice were randomly divided into two groups: MOCK group, and TED group. For the TED group, each mouse was given intramuscular injections of 2 × 10^9^ particles of Ad-TSHR289^36^ in 50 μL PBS (designed by Prof. Ximing Liu, Guang’anmen Hospital, China Academy of Chinese Medical Sciences, produced by DonghuaFang Biological Co., Ltd., Beijing) suspended in 50 μL PBS. The MOCK group received intramuscular injections of an equivalent amount of empty-vector adenovirus in 50 μL PBS. Injections were performed once every three weeks, totaling ten administrations. Throughout the experimental period, the mice were kept in a regulated environment with a 12-hour light/12-hour dark cycle, provided with unrestricted access to food and water.

### Animal treatment

Following model induction, serum levels of T3, T4, and TSH receptor autoantibody (TRAb) were measured in each mouse. The model was considered successfully established when serum thyroid hormone and TRAb concentrations were elevated to at least twice the levels observed in the MOCK group, and MRI revealed significant hypertrophy of the extraocular muscles. Subsequently, the TED mice were randomly divided into three groups and treated daily via intraperitoneal injection for 6 weeks (week 25 to 31) with either PBS (control), sodium lactate (0.2 g/kg, HY-B2227B, MedChemExpress, China), or oxamate (0.75 g/kg, HY-W013032A, MedChemExpress, China). The treatment regimen is illustrated in Figure 2G.

Following TED mouse model construction, the mice were randomly divided into two groups. At week 21, the mice were anesthetized with isoflurane and received periorbital injections of AAV-shNC or AAV-shCREB (2×10^12^ vg/mL) in 25 μL of PBS. The treatment regimen is illustrated in Figure 4F.

### AAV-mediated short hairpin RNA knockdown of CREB in mouse orbit

For in vivo knockdown, AAV vectors encoding CREB-targeting shRNA or a negative control (pAAV-U6-shRNA-CMV-EGFP) were designed and constructed by RIBOBIO (Guangzhou, China). A total of 2 × 10¹² viral particles in 50 μL of PBS were injected locally into the mouse orbit. Transduction efficiency was confirmed via fluorescence microscopy, and CREB knockdown was subsequently validated by western blotting.

### Magnetic resonance imaging

MRI was conducted on a 9.4T BioSpec scanner (Bruker) using a 3-channel surface coil and an 86-mm volume coil. T2-weighted images were acquired using a Turbo RARE sequence (TR/TE = 2500/33 ms; FOV = 2.0 × 2.0 cm²; matrix = 256 × 256; slice thickness = 0.30 mm). To suppress chemical shift artifacts and enable accurate tissue characterization, a fat-water separation method was implemented using a Dixon or iterative decomposition of water and fat with echo asymmetry and least-squares estimation (IDEAL) sequence, which generated separate water-only, fat-only, in-phase, and opposed-phase image sets. Region of interest (ROI) analysis of the MRI images was performed using the Medical Imaging Interaction Toolkit (MITK) software^37^. Briefly, the workflow included importing DICOM image data into MITK, applying N4 bias correction and denoising for image enhancement, and segmenting key orbital structures, including the globe, nerves, extraocular muscles, and adipose tissue, using manual, semi-automated, or atlas-based segmentation techniques. Quantitative analysis was performed for area measurements, volumetric measurements, and signal intensity analysis, with fat-water separation allowing independent evaluation of fat signal contributions within each segmented ROI.

### cAMP assay

Following overnight starvation in 1% FBS, cells seeded in 24-well plates were incubated in antibiotic-free, serum-free medium containing 0.5 mM IBMX to inhibit cAMP decay. The cells were subsequently treated with aptamers, with or without M22 (100 ng/mL, RSR, UK). Intracellular extracts were harvested, stored at -80°C, and analyzed using a competitive polyclonal antibody-based cAMP assay kit (KGE002B, R&D Systems, USA) according to the manufacturer’s instructions and previous descriptions^38^.

## Statistical analysis

Statistical analysis was conducted using GraphPad Prism 10 (GraphPad Software, La Jolla, CA), and the data were presented as the mean ± SEM. The Shapiro-Wilk method was used to determine whether the data were normally distributed, and the homogeneity of variance was tested using the Levene method. If the measurements between two groups were normally distributed, the unpaired Student’s t-test was used; otherwise, the Mann-Whitney U test was used. One-way analysis of variance with post hoc contrasts by the Tukey test was applied to compare the means of multiple groups. A *p* value less than 0.05 was considered statistically significant.

## Acknowledgments

This work was supported by the National Key R&D Program of China (2021YFA1101200 and 2021YFA0909400), the National Natural Science Foundation of China (82171048, 92253201, 32350026, and 22334005), the Ministry and the province of Zhejiang Medical and Health Science and Technology Project (WKJ-ZJ-2336), the Key science and technology program of Wenzhou (ZY2022021), the Fundamental and Interdisciplinary Disciplines Breakthrough Plan of the Ministry of Education of China (JYB2025XDXM602), the Scientific Research Program of FuRong Laboratory (2025PT5013) and the Hunan Science and Technology Innovation Plan (2025ZYJ003). We thank the Scientific Research Center of Wenzhou Medical University for providing excellent consultation and instrumental support.

