## Supplementary for "Immunometabolic Reprogramming by Thyroid-Stimulating Antibodies Drives Orbital Adipogenesis via Histone Lactylation in Thyroid Eye Disease"

### Supplementary materials

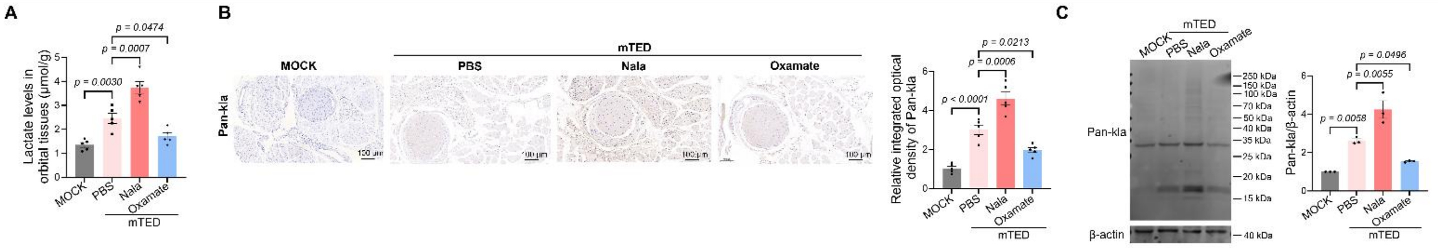


**Supplementary Fig. 1.** Sodium lactate administration increased orbital lactate and pan-lactylation levels in mice, whereas Oxamate administration alleviated these effects. (A) Lactate levels in mouse orbits from the four experimental groups. (B) IHC staining images of Pan-kla in orbital tissues from the four experimental groups and corresponding quantitative analysis of relative integrated optical density (n = 5). Scale bar = 100 μm. (C) The global lactylation levels of orbital tissues in the four experimental groups (n = 3). β-actin served as the loading control. All data are represented as the mean ± SEM. One-way ANOVA, followed by Tukey's multiple post hoc test, was performed in (A-C). Accurate *p*-values are listed in the figures.

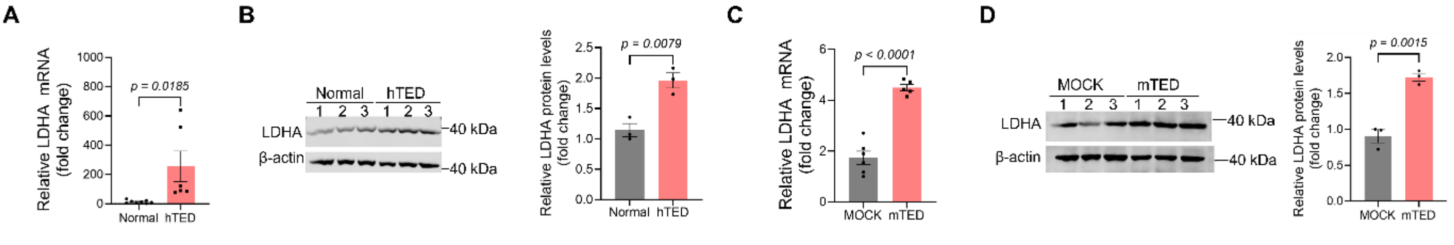


**Supplementary Fig. 2.** LDHA mRNA and protein levels were increased in the orbits from TED patients and TED mice. (A) The mRNA expression levels of LDHA in orbital adipose tissues from normal donors (n = 8) and TED patients (n = 6) by qRT-PCR. (B) Western blot analysis of LDHA in orbital tissues from normal donors (n = 3) and TED patients (n = 3). β-actin served as the loading control. (C) The mRNA expression levels of LDHA in mouse orbital tissues from the MOCK group and TED group by qRT-PCR (n = 5). (D) Western blot analysis of LDHA in mouse orbital tissues from the two mouse groups (n = 3). β-actin served as the loading control. All data are represented as the mean ± SEM. Two-tailed unpaired Student's t-tests were performed in (A-D). Accurate *p*-values are listed in the figures.

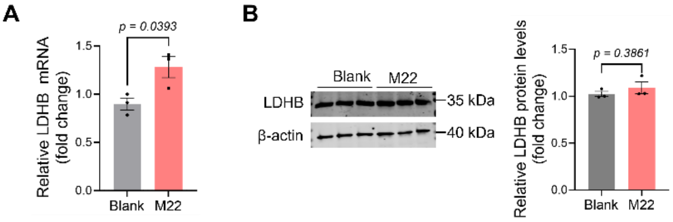


**Supplementary Fig. 3.** M22 treatment had no significant effect on LDHB levels in OFs. (A) The mRNA expression levels of LDHB in OFs stimulated with or without M22 by qRT-PCR (n = 3). (B) Western blot analysis of LDHB expression in OFs stimulated with or without M22 (n = 3). β-actin served as the loading control. All data are represented as the mean ± SEM. Two-tailed unpaired Student's t-tests were performed in (A-B). Accurate *p*-values are listed in the figures.

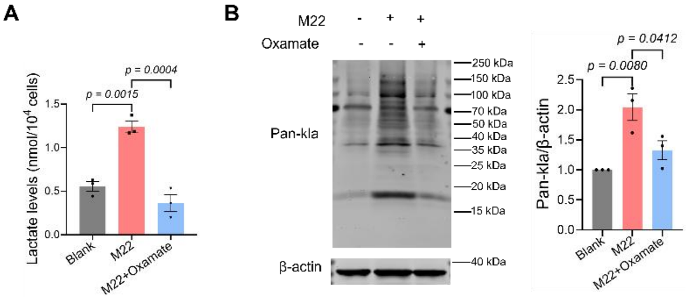


**Supplementary Fig. 4.** Oxamate attenuated M22-induced lactate and pan-lactylation levels. (A) Lactate levels in OFs measured 48 h after M22 stimulation, with or without Oxamate treatment (n = 3). (B) Western blot analysis of global lactylation levels in OFs following 48 h of M22 stimulation, with or without Oxamate treatment (n = 3). β-actin served as the loading control. All data are represented as the mean ± SEM. One-way ANOVA, followed by Tukey's multiple post hoc test, was performed in (A-B). Accurate *p*-values are listed in the figures.

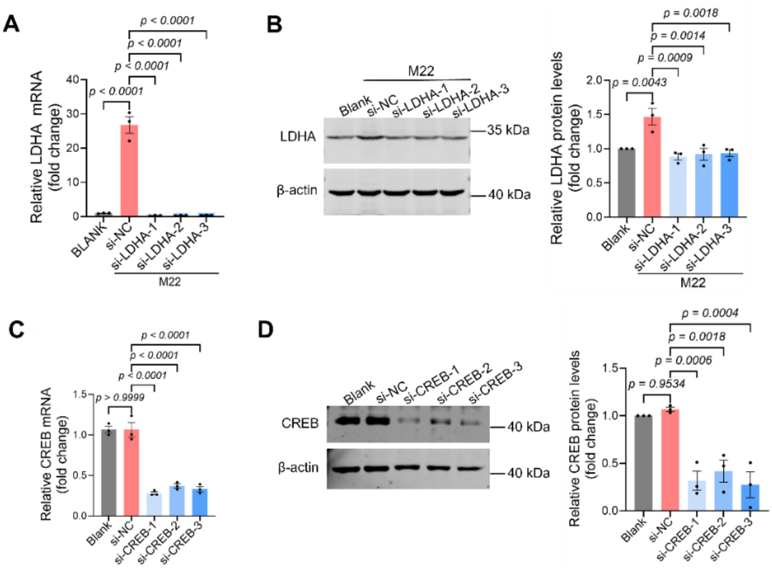


**Supplementary Fig. 5.** si-LDHA and si-CREB effectively reduce LDHA and CREB mRNA and protein levels, respectively. (A) The mRNA expression levels of LDHA in OFs transfected with the indicated siRNAs and stimulated with or without M22 by qRT-PCR (n = 3). (B) Western blot analysis of LDHA in OFs transfected with indicated siRNAs (n = 3). β-actin served as the loading control. (C) The mRNA expression levels of CREB in OFs transfected with the indicated siRNAs by qRT-PCR (n = 3). (D) Western blot analysis of CREB in OFs transfected with indicated siRNAs (n = 3). β-actin served as the loading control. All data are represented as the mean ± SEM. One-way ANOVA, followed by Tukey's multiple post hoc test, was performed in (A-D). Accurate *p*-values are listed in the figures.

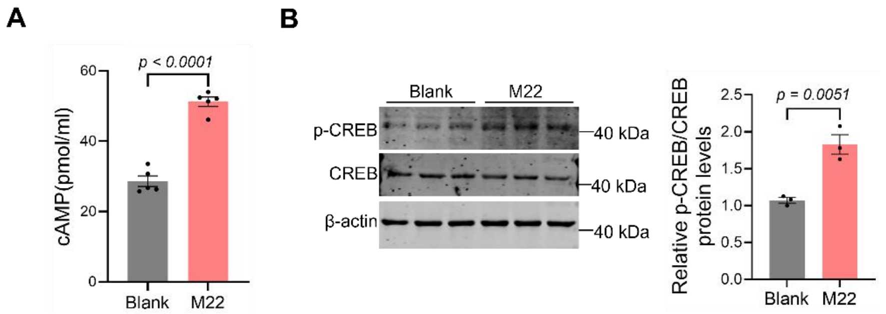


**Supplementary Fig. 6.** M22 activated TSHR via the cAMP-CREB axis. (A) The cAMP levels in OFs stimulated with or without M22 (n = 5). (B) Western blot analysis of p-CREB and CREB in OFs with or without M22 stimulation (n = 3). β-actin served as the loading control. All data are represented as the mean ± SEM. Two-tailed unpaired

Student’s t-tests were performed in (A-B). Accurate *p*-values are listed in the figures.

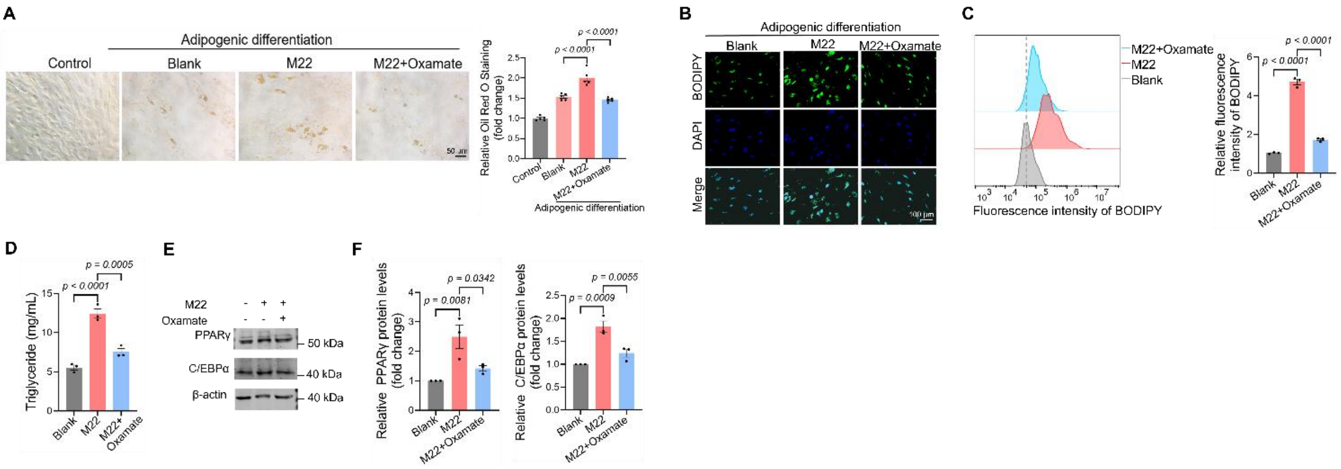


**Supplementary Fig. 7.** Oxamate treatment attenuated M22-induced adipogenesis. (A) Representative Oil Red O staining images of OFs following adipogenic differentiation and M22 or Oxamate treatment (n = 5). Scale bar = 50 µm. (B) Representative BODIPY staining images of adipogenic-differentiated OFs following M22 or Oxamate treatment. (C) Flow cytometry analysis of adipogenic-differentiated OFs following M22 or Oxamate treatment. Scale bar = 100 µm. (D) Triglyceride content in adipogenic-differentiated OFs following M22 or Oxamate treatment (n = 3). (E) Western blot analysis of PPARγ and C/EBPα in adipogenic-differentiated OFs following M22 or Oxamate treatment (n = 3). β-actin served as the loading control. (F) Corresponding quantitative analysis of relative PPARγ and C/EBPα protein levels in panel (E). All data are represented as the mean ± SEM. One-way ANOVA, followed by Tukey's multiple post hoc test, was performed in (A, C-F). Accurate *p*-values are listed in the figures.

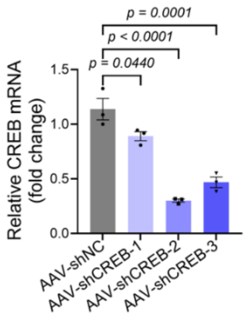


**Supplementary Fig. 8.** The CREB mRNA expression levels in mouse OFs transduced with the indicated AAV-shRNAs, determined by qRT-PCR (n = 3). All data are represented as the mean ± SEM. One-way ANOVA, followed by Tukey's multiple post hoc test, was performed. Accurate *p*-values are listed in the figures.

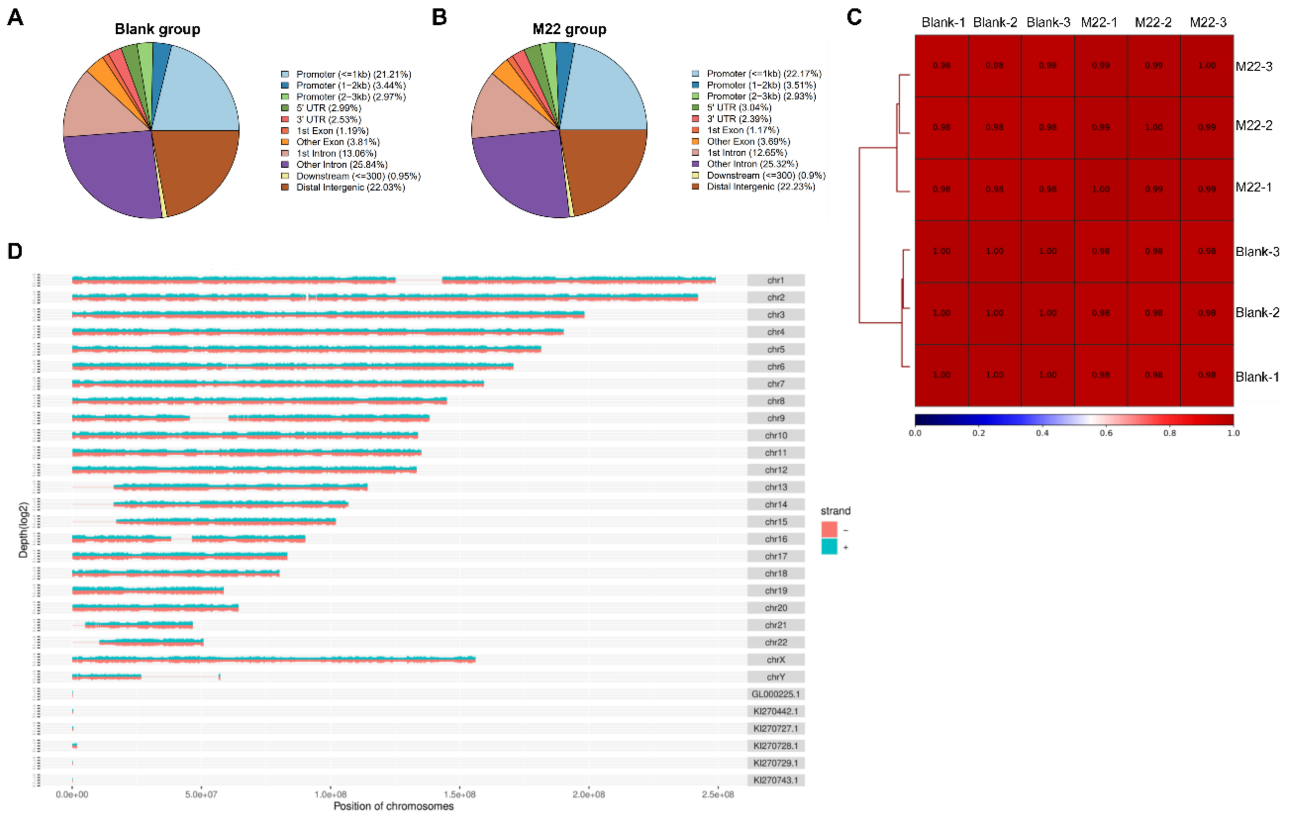


**Supplementary Fig. 9.** CUT&Tag analysis of H3K18la in OFs with or without M22 treatment. (A-B) Genomic distribution of H3K18la in OFs from the Blank and M22 groups. (C) Pearson correlation of fold enrichment between two groups. (D) Location of identified peaks in chromosomes.
